# The role of LmeA, a mycobacterial periplasmic protein, in stabilizing the mannosyltransferase MptA and its product lipomannan under stress

**DOI:** 10.1101/2020.06.18.159426

**Authors:** Kathryn C. Rahlwes, Sarah H. Osman, Yasu S. Morita

## Abstract

The mycobacterial cell envelope has a diderm structure, composed of an outer mycomembrane, an arabinogalactan-peptidoglycan cell wall, periplasm and an inner membrane. Lipomannan (LM) and lipoarabinomannan (LAM) are structural and immunomodulatory components of this cell envelope. LM/LAM biosynthesis involves a number of mannosyltransferases and acyltransferases, and MptA is an α1,6-mannosyltransferase involved in the final extension of the mannan backbones. Recently, we reported the periplasmic protein LmeA being involved in the maturation of the mannan backbone in *Mycobacterium smegmatis*. Here, we examined the role of LmeA under stress conditions. We found that the *lmeA* transcription was upregulated under two stress conditions: stationary growth phase and nutrient starvation. Under both conditions, LAM was decreased, but LM was relatively stable, suggesting that maintaining the cellular level of LM under stress is important. Surprisingly, the protein levels of MptA were decreased in *lmeA* deletion mutant (Δ*lmeA*) in both stress conditions. The transcript levels of *mptA* in Δ*lmeA* were similar to or even higher than those in the wildtype, indicating that the decrease of MptA protein was a post-transcriptional event. Consistent with the decrease in MptA, Δ*lmeA* was unable to maintain the cellular level of LM under stress. Even during active growth, overexpression of LmeA led the cells to produce more LM and become more resistant to several antibiotics. Altogether, our study reveals the roles of LmeA in the homeostasis of the MptA mannosyltransferase particularly under stress conditions, ensuring the stable expression of LM and the maintenance of cell envelope integrity.

---

*Mycobacterium tuberculosis* is the leading cause of death due to a bacterial pathogen, claiming 1.2 million deaths in 2018 alone (1). Furthermore, nearly half a million new cases of multi-drug resistant *M. tuberculosis* were reported (1), indicating urgent needs for discovering novel drug targets. Mycobacteria have one of the most impermeable cell envelopes of bacteria due to its waxy surface, creating an effective barrier against antibiotics and host immune attacks. In a currently proposed model, the outer membrane (OM) is made of the outer leaflet composed of various (glyco)lipids and the inner leaflet primarily composed of mycolic acids. These mycolic acids are linked to arabinan in the arabinogalactan (AG) layer. This layer in turn is covalently attached to the peptidoglycan (PG) cell wall (2–6).

Phosphatidylinositol mannosides (PIMs), lipomannan (LM) and lipoarabinomannan (LAM) are mannose-rich molecules, in which the mannose chain is linked to a phosphatidylinositol (PI) membrane anchor. PIMs are glycolipids carrying up to six mannose residues. They are suggested to be components of the plasma membrane (7, 8), playing both structural and functional roles. For example, lack of hexamannosyl PIM species such as AcPIM6 and Ac_2_PIM6 in *Mycobacterium smegmatis* results in plasma membrane invaginations, a growth delay, an increase in antibiotic sensitivity and a defective permeability barrier (8, 9). LM and LAM are lipoglycans much larger than PIMs, carrying more than 20 mannose residues. These lipoglycans are thought to be present in both the inner membrane and the outer membrane of mycobacteria (2), and appear to play a structural role in cell envelope integrity (10). In *M. tuberculosis*, LAM is important for the effective establishment of infection in mice and interacts with several host receptors, such as Dectin-2, DCSIGN, mannose receptor, and lactosylceramide lipid rafts (11–14). Furthermore, changes in the chain length of LM/LAM decrease virulence of *M. tuberculosis* in mice (10).

The biosynthesis of PIMs/LM/LAM starts with PI. Two mannoses are added from GDP-mannose by sequential actions of two mannosyltransferases, PimA and PimB’, forming PIM2. One or two acyl chains are then added, forming AcPIM2 and Ac_2_PIM2 (15–17). AcPIM2 is further modified by two additional mannoses, which are added by an unknown mannosyltransferase(s), resulting in the formation of AcPIM4. AcPIM4 is the branching point between AcPIM6 and LM/LAM biosynthesis. PimE, an α1,2 mannosyltransferase, adds the fifth mannose from polyprenol phosphate mannose (PPM), committing the pathway to AcPIM6 synthesis (8). AcPIM4 feeds into LM/LAM biosynthesis through an unknown mannosyltransferase, which produces an LM intermediate carrying 5-12 mannose residues. From there, two PPM-dependent mannosyltransferases are involved in producing mature LM, which carries 21-34 mannose residues. MptA, an α1,6 mannosyltransferase, elongates the mannan backbone while MptC, an α1,2 mannosyltransferase, decorates the α1,6 mannan backbone with α1,2 mono-mannose side chains (18–22). The addition of an arabinan chain consisting of ∼70 arabinose residues to LM finally leads to the production of LAM, and a number of arabinosyltransferases are involved in this process (2, 3, 5).

Previously, we have shown that the conserved protein LmeA is involved in the final maturation steps of LM biosynthesis in *M. smegmatis*, potentially aiding MptA in mannan elongation (23). Interestingly, in *M. tuberculosis*, LmeA is predicted to be essential, and upregulated upon mouse infection (24–26). In this study, we examined the role of LmeA in *M. smegmatis* under stress conditions, and demonstrated that LmeA maintains the stable expression of MptA, and its expression level correlates with the cellular LM content and cell envelope integrity.

## RESULTS

### M. tuberculosis *ortholog is functional in* M. smegmatis

LmeA is a conserved protein in Actinobacteria (23). In *M. tuberculosis*, a protein that is 60% homologous to *M. smegmatis* LmeA at the amino acid level is encoded in the gene locus *rv0817c*. The expression of this gene is upregulated in *M. tuberculosis* upon mouse infection (26), implying that LmeA is crucial for the pathogenesis. To validate that Rv0817c is the functional ortholog in *M. tuberculosis*, we cloned the gene and expressed in *M. smegmatis lmeA* deletion mutant (Δ*lmeA*). As shown previously (23), Δ*lmeA* accumulated small LM and diffusely distributed LAM, and *M. smegmatis lmeA* gene complemented the mutant phenotype (Fig. 1A). When the *M. tuberculosis* homolog (*rv0817c*) was introduced to the deletion mutant instead of *M. smegmatis lmeA*, it was able to restore the LM/LAM phenotype of *M. smegmatis* Δ*lmeA* (Fig. 1A). LmeA does not have an impact on the biosynthesis of structurally related PIM species, and as expected, *M. tuberculosis* LmeA showed no effect on their biosynthesis (Fig. 1B). These data suggested that Rv0817c is functionally equivalent to *M. smegmatis* LmeA. Therefore, *M. tuberculosis* LmeA, which has been reported to be critical for growth and upregulated upon mouse infection (24–26), is a functional ortholog, suggesting its involvement in mannan elongation in *M. tuberculosis*.

**Figure 1.**
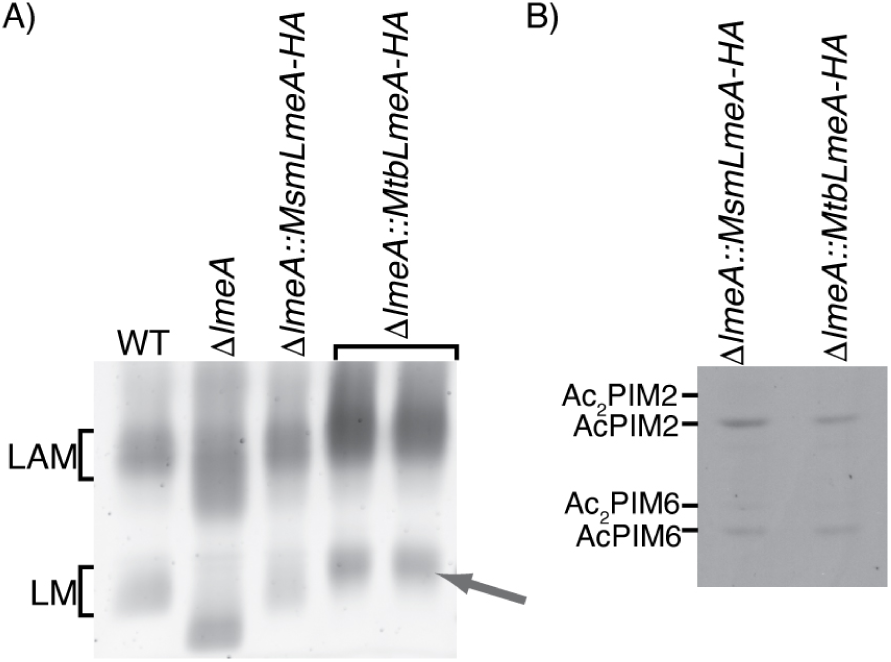
*M. tuberculosis* LmeA is the functional ortholog of *M. smegmatis* LmeA. The Δ*lmeA M. smegmatis* was complemented with the *M. tuberculosis* LmeA ortholog (*rv0817c*) expressed at mycobacteriophage L5 attB site. A) SDS-PAGE analysis of LM and LAM. Two independent clones of Δ*lmeA*::*Mtb_lmeA* were able to restore mature LM formation (arrow). B) HPTLC analysis of PIMs. PIM profiles were not affected by complementation of *Mtb_lmeA-HA*.

### M. smegmatis lmeA *is upregulated upon stress*

The upregulation of *M. tuberculosis lmeA* (*rv0817c*) during mouse infection implies that expression of this gene may be a response to environmental stress. Given that Rv0817c is orthologous to *M. smegmatis* LmeA, we speculated that LmeA may respond to stress conditions in *M. smegmatis* as well. To examine if *lmeA* gene expression is upregulated upon stress, we extracted RNA from cells under stress and examined the transcript levels of *lmeA* in *M. smegmatis* strains (wildtype (WT), Δ*lmeA*, and Δ*lmeA*::P_native_*lmeA-HA*) by quantitative RT-PCR (qRT-PCR). As a model stress condition, we first tested the stationary growth phase and found that *lmeA* was induced 2.5 fold by 72 h in WT cells when compared to log (Fig. 2A). In addition, Δ*lmeA* mutant complemented with *lmeA* gene, which carries its upstream 165 bp as a putative endogenous promoter region (Δ*lmeA*::P_native_*lmeAHA*)(23), showed a similar trend of upregulation although it was not statistically significant.

**Figure 2.**
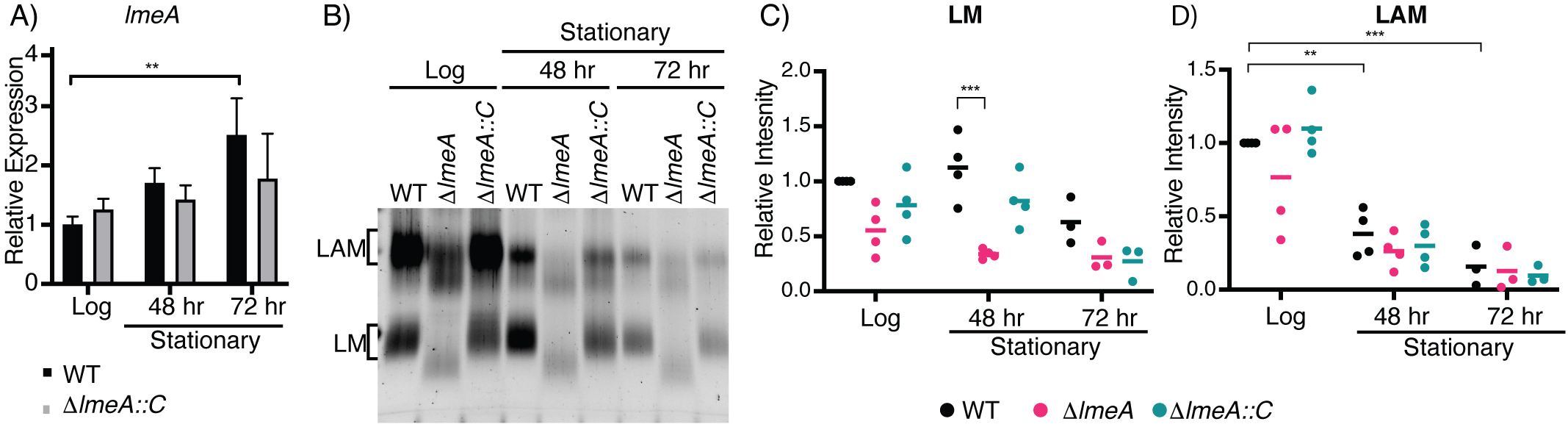
Expression of *lmeA* transcript and cellular levels of LM/LAM during stationary growth phase. A) Relative expression of *lmeA* transcript normalized to a housekeeping gene *gyrB*. A significant increase in *lmeA* expression is noted by 72 h of growth in comparison to log (18 h). **, p < 0.05 by ANOVA. B) SDS-PAGE analysis of LM/LAM during stationary phase. A representative image of biological triplicate is shown. LAM decreased significantly more in the stationary phase than LM. C-D) Dot plots showing intensities of LM (C) and LAM (D) from biological replicates. Signal intensities were quantified using ImageQuant and normalized to the intensity of WT at log phase. The averages are shown as horizontal bars. **, p < 0.01; *, p < 0.05 by ANOVA. Δ*lmeA*::*C*, Δ*lmeA*::P_native_*lmeA-HA* (also see text).

The cellular levels of LAM rapidly decline in stationary phase, but the levels of LM remain relatively constant regardless of the growth phase (21, 27). Because LmeA plays a role in mannan biosynthesis (23), we speculated that the deletion of *lmeA* may affect the LM maintenance during stationary phase. Consistent with previous studies, we noted a 2.5-fold decrease in LAM levels in WT by the 72 h time point, and similar declines in LAM levels were observed in Δ*lmeA* and Δ*lmeA*::P_native_*lmeA-HA* in the stationary phase (Fig. 2B, 2D, S1A and S1B). In contrast, the LM levels of WT cells were not significantly different between log and stationary phases (Fig. 2B, 2C, S1A and S1B). Strikingly, LM levels in the Δ*lmeA* mutant were 3-fold lower than those in WT at the 48 h time point (Fig. 2C). The observed decline in LM levels in the Δ*lmeA* mutant is specific, as there were no specific changes in the levels of other lipids such as phospholipids and PIMs that can be attributable to the lack of LmeA (Fig. S1C). Overall, loss of LmeA not only makes mannan elongation defective, but also impacts the maintenance of LM levels during stationary phase.

### LmeA is critical for the maintenance of MptA during stress

Since LmeA is involved in the biosynthetic step of LM at or near the MptA-mediated mannan elongation (23), we speculated that the lack of LmeA may affect MptA abundance, leading to the decreased amount of LM during stationary phase. To examine this possibility, we examined the MptA proteins levels through anti-MptA western blots at each time point (Fig. 3A). The abundance of MptA was maintained at similar levels during the stationary phase in WT cells. In contrast, in Δ*lmeA*, the protein level was reduced at both 48 and 72 h time points and these reductions in MptA abundance were alleviated in the complemented strain. To confirm that this decrease is not a general reduction in protein abundance, we examined changes in the abundance of another mannosyltransferase, MptC, which mediates the α1,2 monomannose addition. The levels of MptC were comparable among the three strains. There were no specific changes in its abundance that can be attributed to the *lmeA* gene deletion (Fig. 3B). We therefore conclude that LmeA is necessary for the maintenance of MptA protein.

**Figure 3.**
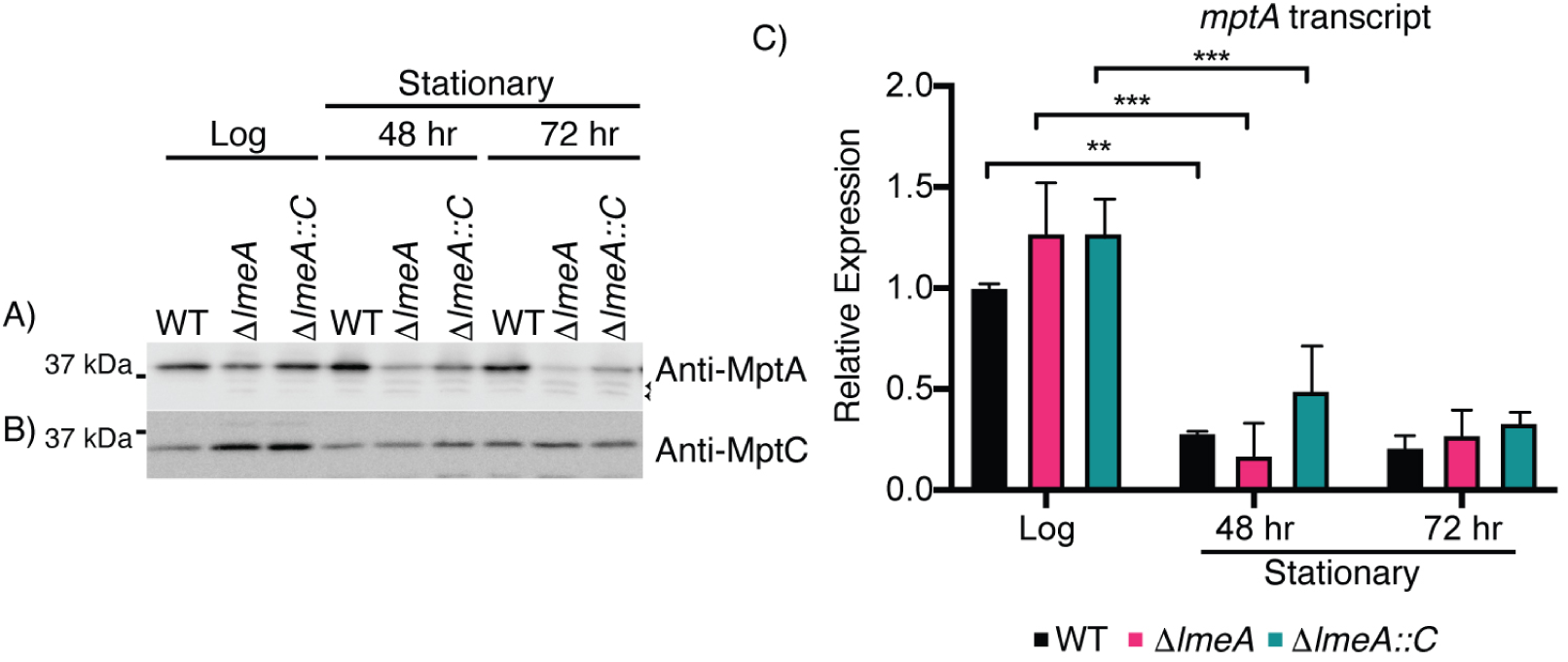
Protein levels of MptA decrease significantly in Δ*lmeA* during stationary phase. A-B) Western blot analysis of MptA (A) and MptC (B) over a 72 h time course of growth. MptA shows significant decrease in Δ*lmeA*. Arrowheads, potential degradation products of MptA. MptC remains fairly constant between WT, Δ*lmeA*, and Δ*lmeA*::P_native_*lmeA-HA.* For SDS-PAGE, an equal concentration of protein was loaded to each lane. C) Relative expression levels of *mptA* transcript was normalized to *gyrB*. A dramatic decrease in mRNA transcript levels is noted between log and 48 h stationary phase. ***, p < 0.001; **, p < 0.05 by ANOVA. There is no significant differences among WT, Δ*lmeA* and Δ*lmeA*::P_native_*lmeA-HA* at all time points. Δ*lmeA*::*C*, Δ*lmeA*::P_native_*lmeA-HA.*

Is the reduced abundance of MptA protein in Δ*lmeA* stationary phase cells due to downregulation of *mptA* transcription in the mutant? To answer this question, we determined the relative levels of *mptA* transcript by qRT-PCR. While there were no significant differences in the expression of *mptA* between the three strains, a general decrease was observed in *mptA* transcript levels at 48 h in comparison to the log phase cells (Fig. 3C). After this time point, the transcript levels of *mptA* were stable with no further decrease. The result that mRNA levels were comparable between WT and Δl*meA* suggests that MptA is degraded at the protein level during stress conditions in the absence of LmeA.

Next, we used starvation as another condition to characterize stress on LM/LAM biosynthesis. We starved *M. smegmatis* strains (WT, Δ*lmeA*, and Δ*lmeA*::P_native_*lmeA-HA*) in phosphate-buffered saline (PBS) for 24 h. We found increase in *lmeA* transcripts upon starvation, which was also observed in Δ*lmeA*::P_native_*lmeA-HA* (Fig. S2A). Consistent with the trend of transcriptional upregulation in the complemented strain, we detected a higher level of LmeA-HA protein when Δ*lmeA*::P_native_*lmeA-HA* was starved (Fig. S2B). As observed in stationary phase cells, there was a substantial depletion of LAM upon starvation while LM remained relatively constant in WT (Fig. 4A, S3A and S3B). Similar to the observed depletion of LM in stationary phase Δ*lmeA*, LM abundance in Δ*lmeA* during starvation was at ∼50% level of that in WT. Other membrane phospholipids appear similar among the three strains (Fig. S3C). Furthermore, MptA protein levels were ∼20 times less in Δ*lmeA* than in WT: a more dramatic fold change than that during stationary phase. In contrast, changes in MptC levels were much milder, being less than 2-fold (Fig. 4B and C). The Δ*lmeA*::P_native_*lmeA* was able to partially complement the phenotype, although MptA levels were 3.5-fold less than that in WT. Overall, starvation is another stress condition, where *lmeA* gene expression is upregulated, and the presence of LmeA is critical for the maintenance of MptA and LM.

**Figure 4.**
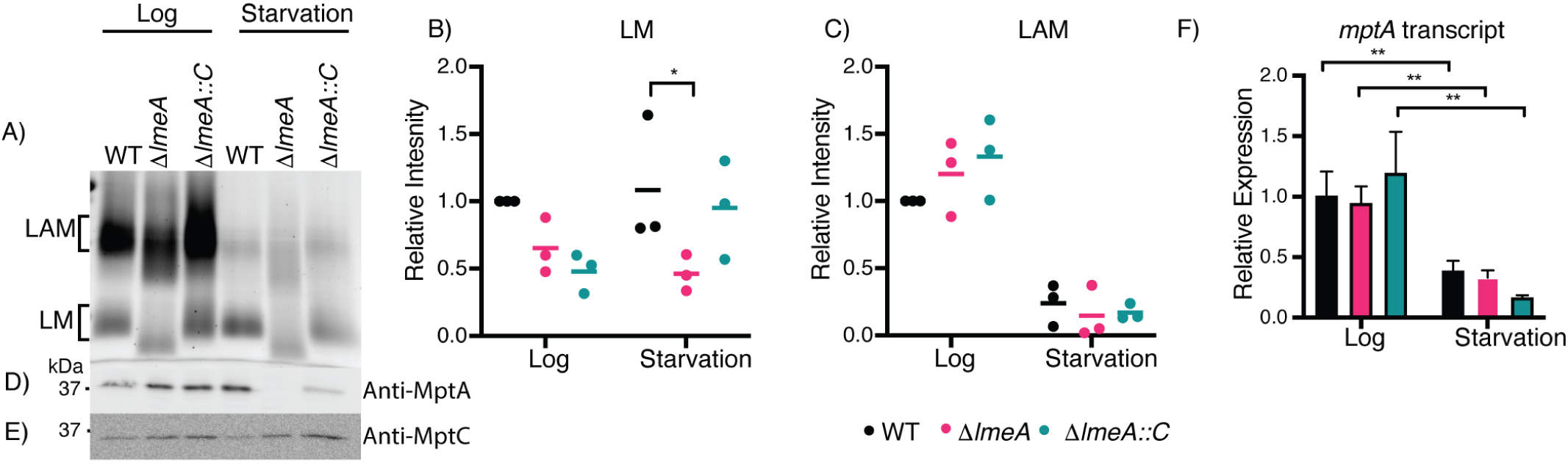
Effect of starvation on cellular levels of LM/LAM and their biosynthetic enzymes. A-C) SDS-PAGE analysis of LM/LAM. LAM decreases during starvation, while LM remains constant. A representative image of biological triplicate is shown. LM (B) and LAM (C) from biological triplicate were quantified and changes relative to WT levels in log phase are shown in dot plot. Average is shown as a horizontal bar. D) Western blot analysis of MptA in log phase and after starvation. MptA is decreased in Δ*lmeA* in comparison to WT during starvation. E) Western blot of MptC in log phase and after starvation. No significant decrease was noted among WT, Δ*lmeA*, and Δ*lmeA*::P_native_*lmeA-HA* before and after starvation. An equal amount of proteins was loaded for western blot analysis. F) Transcript levels of *mptA* relative to WT in log phase, normalized to *gyrB*. The levels of *mptA* transcript decreased from log to starvation, but deletion of *lmeA* gene had no apparent impact. **, p < 0.01 by ANOVA. Δ*lmeA*::*C*, Δ*lmeA*::P_native_*lmeA-HA.*

Given that *lmeA* transcription was upregulated upon stress exposure and the deletion of *lmeA* resulted in decreased abundance of LM in both stationary phase and starvation, we considered the possibility that the overexpression of LmeA may impact LM abundance even during active growth. We created an episomal expression vector for LmeA-HA, in which the gene expression is driven by the strong heat shock protein 60 (HSP60) promoter (28). In this LmeA over-expression strain (LmeA OE), LmeA-HA protein levels were ∼10 fold higher than those in Δ*lmeA*::P_native_*lmeA-HA*, where *lmeA-HA* transcription is driven by the putative endogenous promoter (Fig. 5A). In response to the over-expression of LmeA-HA, MptA levels show a slight increase of ∼40% in LmeA OE when compared to WT (Fig. 5B and 5C). Concomitant with the increase in MptA, LM levels were increased by ∼80% (Fig. 5D, 5E, S4A and S4B). In contrast, in Δ*lmeA*, the MptA levels decreased by ∼1.9 fold when compared to WT even in the log phase. The impact of LmeA OE is specific to LM as it does not show significant impact on LAM levels and does not impact the biosynthesis of PIMs and other phospholipids (Fig. S4C). Taken together, these data suggest that LmeA not only prevents the degradation of MptA under stress conditions but also helps to enhance the production of LM during active growth.

**Figure 5.**
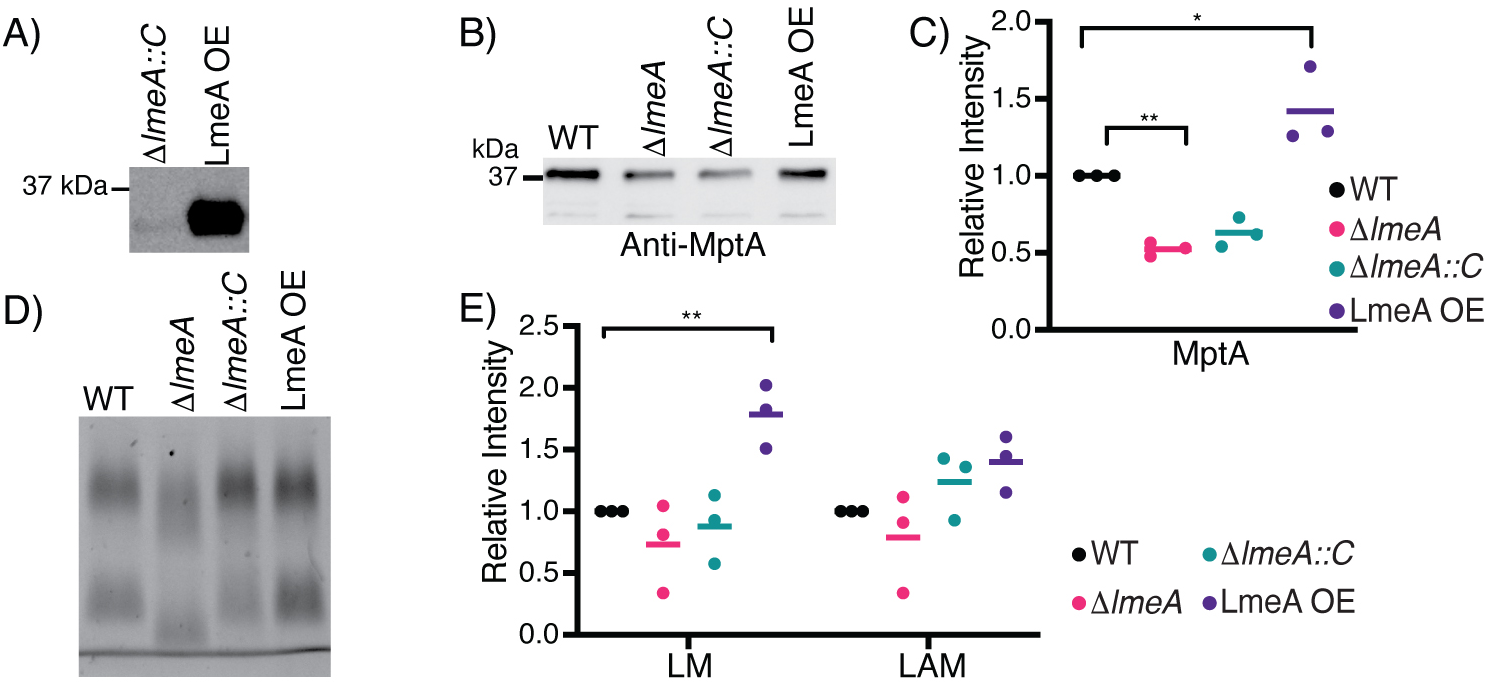
Impact of overexpression of LmeA on LM/LAM biosynthesis. A) Anti-HA western blot to detect LmeA-HA expressed in Δ*lmeA*::P_native_*lmeA-HA* (indicated in the figure as Δ*lmeA*::*C*) and LmeA-HA overexpression strain (LmeA OE). LmeA-HA was ∼10 fold more intense in LmeA OE than in Δ*lmeA*::P_na-_ tive*lmeA-HA*. B) Western blot analysis of MptA in actively growing WT, Δ*lmeA*, Δ*lmeA*::P_native_*lmeA-HA*, and LmeA-HA OE. An equal amount of proteins was loaded to each lane. A representative image of biological triplicate is shown. Δ*lmeA*::*C*, Δ*lmeA*::P_native_*lmeA-HA.* C) Dot plot showing relative abundance of MptA in biological triplicate. Intensity was normalized to WT and the average is shown as a horizontal bar. *, p < 0.05; **, p < 0.01 by ANOVA. D) Cellular levels of LM and LAM in actively growing WT, Δ*lmeA*, Δ*lmeA*::P_native_*lmeA-HA*, and LmeA OE. A representative image of three biological replicates is shown. E) Dot plot of cellular LM/LAM levels. The average of biological replicates is shown as a horizontal bar. **, p < 0.01 by ANOVA.

### The expression level of LmeA correlates with antibiotic resistance

Previously, using deletion mutants of *mptA* and *mptC*, LM/LAM have been shown to play important roles in conferring intrinsic resistance to various antibiotics (10, 21). Given the significant impacts of *lmeA* deletion and over-expression on LM/LAM as shown above and previously (23), we examined the impact of these changes in LM/LAM on antibiotic resistance. We determined the IC_90_ of five antibiotics that target either periplasmic peptidoglycan biosynthesis (ampicillin, cefotaxime, vancomycin) or cytoplasmic protein synthesis (clarithromycin and erythromycin) (Table 1). The Δ*lmeA* mutant was more sensitive to all five antibiotics relative to WT. In particular, IC_90_ for cefotaxime and erythromycin were >10 and 8-fold reduced in Δ*lmeA* in comparison to WT, respectively. The complemented strain (Δ*lmeA*::P_native_*lmeA*) restored resistance to all antibiotics, but was only partial restoration for vancomycin, cefotaxime and erythromycin. As we demonstrated previously that antibiotic sensitivity is dependent on growth medium (9), we further tested sensitivities to these antibiotics in a medium different from Middlebrook 7H9, our standard growth medium. As shown in Table 2, WT *M. smegmatis* grown in M63 medium became notably more resistant to vancomycin than those grown in Middlebrook 7H9. IC_90_ for clarithromycin and erythromycin also became higher in M63 than in Middlebrook 7H9. Nevertheless, the increased sensitivities to antibiotics found in Δ*lmeA* were reproducible in M63, and the complemented strain restored antibiotic resistance at least partially (Table 2). Taken together, Δ*lmeA* is more sensitive to various antibiotics, likely due to truncated mannan backbone of LM and LAM compromising the cell envelope integrity.

**Table 1.** Antibiotic sensitivity of *M. smegmatis* strains grown in Middlebrook 7H9. IC_90_ (µg/mL) is shown as average ± standard deviation.

|  | Vancomycin | Cefotaxime | Ampicillin | Clarithromycin | Erythromycin |
| --- | --- | --- | --- | --- | --- |
| **WT** | 1.00 ± 0.14 | >100 | >100 | 0.15 ± 0.01 | 0.96 ± 0.17 |
| ***ΔlmeA*** | 0.39 ± 0.04 | 9.77 ± 2.08 | 70.0 ± 14.8 | 0.05 ± 0.01 | 0.12 ± 0.01 |
| ***ΔlmeA::P<sub>native</sub>lmeA-HA*** | 0.58 ± 0.09 | 48.2 ± 7.6 | >100 | 0.15 ± 0.01 | 0.31 ± 0.01 |
| **LmeA OE** | 2.13 ± 0.33 | >100 | >100 | 0.25 ± 0.05 | 0.92 ± 0.21 |

**Table 2.** Antibiotic sensitivity of *M. smegmatis* strains grown in M63 medium. IC_90_ (µg/mL) is shown as average ± standard deviation.

|  | Vancomycin | Cefotaxime | Ampicillin | Clarithromycin | Erythromycin |
| --- | --- | --- | --- | --- | --- |
| **WT** | >100 | >100 | >100 | 1.61 ± 0.24 | 78.9 ± 28.2 |
| ***ΔlmeA*** | 14.7 ± 2.9 | >100 | >100 | 0.35 ± 0.03 | 0.41 ± 0.16 |
| ***ΔlmeA::P<sub>native</sub>lmeA-HA*** | 99.6 ± 17.9 | >100 | >100 | 0.61 ± 0.02 | 9.51 ± 1.64 |

LmeA OE, the strain overexpressing LmeA in the WT background, showed enhanced LM abundance compared with WT. We wondered if the increased abundance of LM might impact the antibiotic resistance. As shown in Table 1, LmeA OE showed up to ∼2-fold increase in resistance to vancomycin and clarithromycin (Table 1), indicating that the amount of LM correlates with antibiotic resistance.

## DISCUSSION

In this study, we examined the impact of LmeA expression on LM/LAM biosynthesis in detail and demonstrated that the expression of LmeA is critical for MptA stability under stress conditions (Fig. 6). Stability of MptA during stress exposure appears to be important specifically for maintaining LM but not LAM because LAM levels quickly declined during stress exposure irrespective of the presence or absence of LmeA. These observations strongly support that LmeA has a regulatory role in stress response, preventing MptA degradation and supporting LM biosynthesis.

**Figure 6.**
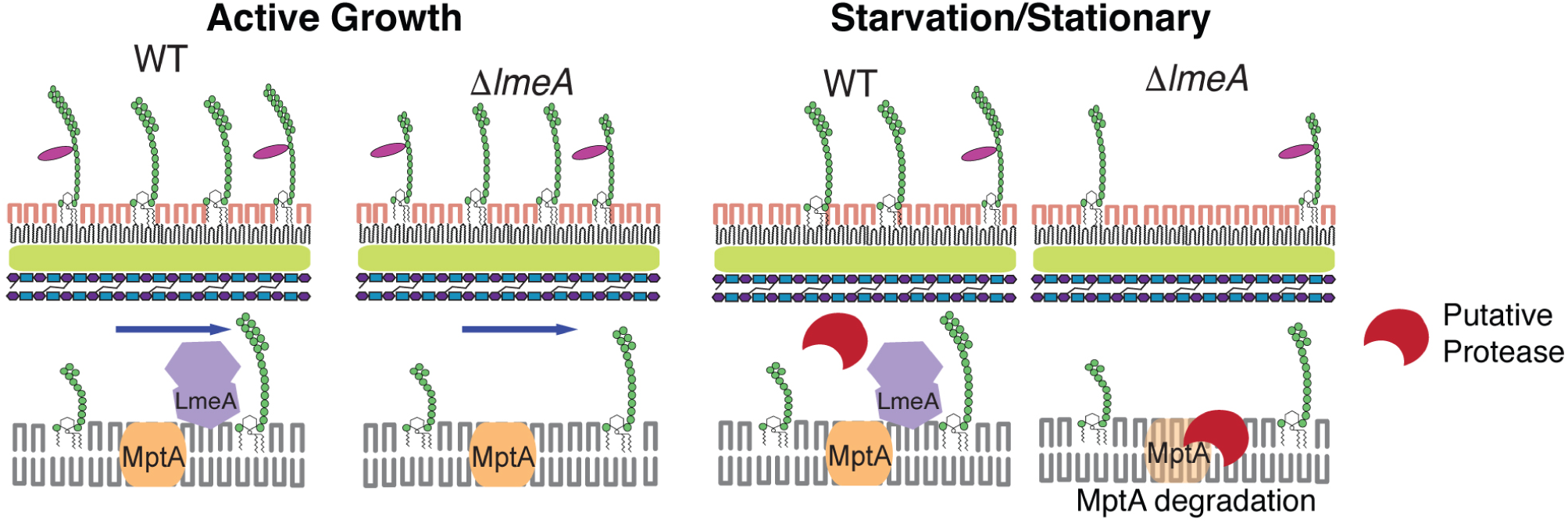
Proposed model of how LmeA functions during stress conditions. Under active growth conditions, LmeA aids MptA in its function, potentially through either stabilizing MptA or facilitating the trafficking of LM/LAM or both. Upon stress exposure such as starvation or stationary phase, LmeA protects MptA from a putative periplasmic protease.

LmeA drives LM biosynthesis by providing stability to the mannosyltransferase MptA. LmeA plays this role at the post-transcriptional level since there is no difference in mRNA levels of *mptA* between Δ*lmeA* and WT. We suggest that the decreased levels of MptA in Δ*lmeA* are mainly due to protein degradation. In fact, two fainter western blot bands migrating below MptA are barely visible in actively growing WT cells but become more prominent under stress conditions (see Fig. 3A), possibly indicating the presence of partially degraded MptA. There are a number of proteases that have the active site in the periplasmic space, including the serine proteases MarP, HtrA and Rv3671c that are proposed to play roles in cell envelope homeostasis and stress response in mycobacteria (29–33). It remains to be determined if MptA is proteolytically degraded and which protease fulfills the role.

How does LmeA prevent the degradation of MptA during stress conditions? LmeA is a distant homolog of bactericidal/permeability increasing protein (BPI) (23), and like BPI, it has a tubular lipid binding (TULIP) lipid transfer domain (34). Furthermore, LmeA also carries a Chorein-N domain, which is also involved in lipid transfer (34). Indeed, we have previously shown that *M. smegmatis* LmeA can bind to phospholipids (23). We speculate that full elongation of mannan chain is coupled with the transfer of LM across the periplasmic space. Such a concurrent process of mannan elongation and lipid transfer would explain why LM intermediates with a short mannan chain accumulate in Δ*lmeA*. We further speculate that, in the absence of LmeA, MptA may be stalled with partially elongated LM intermediates and may be subjected to degradation particularly during stress conditions.

The role of LmeA under stress exposure may be further inferred by the location of the gene in the genome. The gene encoding LmeA appears to form an operon with a downstream gene encoding a putative thioredoxin, ThiX (MSMEG_5786 in *M. smegmatis*, and Rv0816c in *M. tuberculosis* H37Rv), and this operon structure is widely conserved in mycobacteria. ThiX has a predicted signal peptide, suggesting that it is a periplasmic thioredoxin. In Gram-negative bacteria, periplasmic thioredoxins are critical for withstanding redox stress (35). Furthermore, the upstream of *lmeA* is *glnR* (MSMEG_5784 / Rv0818), which is involved in responding to the environmental stress such as redox stress and nitrogen starvation (36, 37). *glnR* forms an apparent operon with *mshD*, the final enzyme for the synthesis of mycothiol (38), which is a key molecule to regulate the reduced environment in the cytoplasm. These surrounding genes suggest that *lmeA* is situated in a potential regulon for the redox stress response. How LM/LAM might be important for resisting redox stress remains unclear. However, it is documented that cell envelope rearrangement is critical for mycobacterial cells when shifting from an actively growing phase to a more stressful non-growing phase (4). LmeA may play an integral role in regulating the cell envelope rearrangement through facilitating the action of the MptA mannosyltransferase.

We showed that LAM rapidly depletes under stress conditions whereas the abundance of LM does not change significantly. These differences in stress response suggest that LM and LAM are playing functionally different roles during stress condition. It remains to be examined if LAM is degraded or shed from the cell under these stress conditions. In either case, LM may play a more vital role in stabilization of the cell envelope under stress conditions. Indeed, LmeA OE showed increase in LM but not LAM (Fig 5 B), and this increase in LM was correlated with increased antibiotic resistance (Table 1). Rather subtle changes in the structure and abundance of LM and LAM have significant impact on the cell envelope integrity. We propose that LmeA is a stress response protein that plays a critical role in the biosynthesis and trafficking of LM/LAM, thereby impacting the cell envelope integrity and antibiotic resistance.

## EXPERIMENTAL PROCEDURES

### Growth conditions

*Mycobacterium smegmatis* mc^2^155 (WT), Δ*lmeA*, Δ*lmeA*::P_native_*lmeA-HA*, Δ*lmeA*::P_hsp60_*lmeA-HA* (23), Δ*lmeA*::*Mtb_lmeA-HA* (this study), and LmeA OE (this study) were grown in a liquid medium of Middlebrook 7H9 base (BD), supplemented with glycerol (0.2%, v/v), glucose (0.2%, w/v), NaCl (15 mM), and Tween80 (0.05%, v/v), at 30°C with shaking at 130 rpm. For analysis, 25 OD_600_ units of cells were centrifuged for 10 min at 3,220x *g* and pellets were processed for bead-beating, lipid extraction, or RNA extraction.

For the starvation condition, the *M. smegmatis* strains were grown in Middlebrook 7H9 at 30°C with shaking at 130 rpm until a logarithmic growth phase (OD_600_ = 0.6-1.0). The cells were then centrifuged for 10 min at 3,220x *g*, resuspended in PBS, and incubated for additional 24 h at the same condition.

### Expression vectors

Two vectors, Mtb_LmeA expression vector (pMUM105) and LmeA overexpression vector (pMUM063), were constructed as detailed below.

### pMUM063

*lmeA* was excised from pMUM054 (23) by digestion with Van91I and XbaI and ligated into pYAB017 (39), which was digested with Van91I, XbaI, and SphI. pYAB017 has an *oriM* and does not integrate into the mycobacterial chromosome. After ligation, the plasmid was electroporated into WT *M. smegmatis* to create the LmeA overexpression strain.

### pMUM105

The gene encoding Rv0817c was PCR-amplified from *M. tuberculosis* genomic DNA using primers A367 and A368 (Table S1). The fragment was then ligated into pDO23A (40) through Gateway BP cloning, creating plasmid pMUM089. The attB-Rv0817c region of this plasmid was then amplified with primers A444 and A445. The PCR fragment was then inserted into pMUM082, which was linearized by ClaI and NdeI, through Gibson assembly. The final plasmid construct, pMUM105, was electroporated into *M. smegmatis* Δ*lmeA.*

### Lipids/LM/LAM extraction

Lipids were extracted as described previously (23, 41). Briefly, lipids were extracted from cell pellets by chloroform/methanol/water and purified by *n*-butanol/water phase partition. Phospholipids and PIMs were analyzed by high performance thin layer chromatography (HPTLC) (silica gel 60, EMD Merck) using chloroform/methanol/13 M ammonia/ 1 M ammonium acetate/ water (180:140:9:9:23) as the mobile phase. Phospholipids were visualized by iodine, and PIMs were visualized by orcinol as described (23, 41). LM and LAM were extracted from the delipidated pellets, separated by SDS-PAGE (15% gel), and visualized using ProQ Emerald 488 glycan staining kit (Life Technologies) as described (23, 41). Intensities of LM and LAM were determined using ImageQuant. Statistical significance was determined by two-way ANOVA using Graphad Prism version 8.

### Cell lysis and protein analysis

Cell pellets were lysed by bead-beating as described (23). The protein concentration of each lysate was determined by a BCA assay kit (ThermoFisher Scientific), and adjusted to 1.2 mg/mL. The lysate (12 µL) was mixed with reducing sample loading buffer, denatured on ice for 30 min, loaded onto a 12% SDS-PAGE gel, and electrophoresed at 150 V for 70 min. Western blot was performed as previously described (21) using rabbit anti-MptA, rabbit anti-MptC, or mouse anti-HA (Millipore-Sigma) at 1:2000 dilution as a primary antibody. After incubating with horseradish peroxidase-conjugated anti-rabbit or anti-mouse IgG as a secondary antibody at 1:2000 dilution (GE Healthcare.), protein bands were visualized by chemiluminescence and the image was captured by ImageQuant LAS 4000 mini (GE Healthcare).

### Purification of RNA

RNA was extracted and purified from the cell lysate as described (42). Briefly, cell culture was centrifuged at 3,220x *g* for 10 min. The wet pellets (∼400 mg) were then frozen at -80°C overnight and then resuspend in 1 mL of TRIzol (ThermoFisher Scientific). Cells were lysed with 0.5 mL of zirconia beads using a BeadBug Microtube Homogenizer (Benchmark Scientific) at 4°C with beating at 4,000 rpm for 1 min followed by resting on ice for 2 min. The bead homogenization was performed three times in total and followed by a 5 min incubation at room temperature. The samples were then centrifuged for 30 s at 10,000x *g* at 4°C. The purified RNA was then concentrated using an RNA Clean and Concentrator kit following manufacturer’s protocol (Zymo Research). Two µg of RNA was used to synthesize cDNA using Maxima H minus reverse transcriptase (ThermoFisher Scientific) and 1 µM of random hexamers (Promega) in accordance with manufacturer’s instructions.

### Quantitative PCR

For each 25-µl reaction, 12.5 µL of the Roche FastStart Universal SYBR Green Master mix (Millipore Sigma) was mixed with cDNA and primers to achieve the final concentrations of 100 ng/µl and 400 nM, respectively (Table S1). The quantitative PCR was performed on Stratagene Mx3005P (Agilent Technologies) with the following program: 95°C for 10 min, followed by 40 cycles of incubations at 95°C for 30 s, at 60°C for 30 s, and 72°C for 30 s 72°C. Data were analyzed to determine 2^-^ ΔΔCT using the MxPro software and evaluated for statistical significance through GraphPad Prism version 8.

### Antibiotic sensitivity

Frozen aliquots of WT, Δ*lmeA*, Δ*lmeA*::P_native_*lmeAHA*, Δ*lmeA*::P_hsp60_*lmeA-HA* and LmeA OE were prepared from log phase cultures (OD_600_ = 0.5 – 1.0) grown in Middlebrook 7H9 and the colony forming units (cfu) were determined. In 96-well microtiter plates, antibiotics were serially diluted in 100 µl of either 7H9 or M63 and mixed with 100 µl of cells from the frozen stocks to achieve the final density of 5.0 × 10^3^ cfu/mL. The plates were incubated in a humidity chamber at 37°C for Middle-brook 7H9 or 30°C for M63. After a 24-hour incubation, 20 µL of 0.015% (w/v) resazurin solution was added to initiate colorization. After an additional 8 hour for Middlebrook 7H9 or 13.5 hour for M63 incubation, the plates were read on a spectrophotometer at 570 and 600 nm. Percent difference in cell viability between antibiotic-treated and control cells was calculated as before (9). The IC_90_ values were calculated using OriginPro 9.1 data analysis software.

## Acknowledgement

This work was supported by a Biomedical Research Grant (RG-414805) from the American Lung Association and a Research Grant from the Pittsfield Anti-Tuberculosis Association to YSM. We thank Dr. Sandra Petersen and Kelsey Clements for help with qRT-PCR. We also thank Ian Sparks, Dr. Whitney Blocker McTigue, Dr. Takehiro Kado, and Dr. Breanna Pasko for discussion and critical reading of the manuscript.

